# Midazolam suppresses glioma progression by attenuating neuronal activity and downregulating IGF1 signaling

**DOI:** 10.64898/2026.03.31.715727

**Authors:** Zhihui Qi, Zhiming Ye, Kahei Chan, Yumin Wu, Yu Yu, Yang Hu, Yi Lu, Jiaoyan Ren, Maojin Yao, Zhongxing Wang

## Abstract

Glioma is the most common primary malignant tumor of the brain, and accumulating evidence indicates that neuronal activity plays a pivotal role in tumor progression. In this study, neuronal activity is modulated *in vitro* using potassium chloride (KCl)-induced depolarization and midazolam (MDZ)-mediated suppression. MDZ is a neuronal activity modulation medication, commonly used for sedation, anxiolysis, and amnesia in clinics. After treatment, conditioned media derived from these neuronal cultures are subsequently co-cultured with glioma cells. EdU incorporation assays demonstrate that MDZ significantly inhibits glioma cell proliferation *in vitro*. Furthermore, an orthotopic xenograft glioma model is established to assess the anti-tumor efficacy of MDZ *in vivo*, as evaluated by tumor volume and Ki-67 immunostaining. Mechanistically, insulin-like growth factor 1 (IGF1) is identified as the neuronal-activity-regulated factor that promotes glioma growth through activation of the PI3K/AKT signaling pathway. Moreover, transcriptomic profiling of brain tissues reveals that MDZ attenuates neuronal activity and downregulates neuron-derived growth factors in both glioma and non-tumor regions, thereby exerting anti-tumor effects *in vivo*. Collectively, these findings demonstrate that MDZ suppresses glioma progression by suppressing neuronal activity and inhibiting neuron-derived trophic factors, providing new insights into the development of therapeutic strategies for glioma.

## Introduction

Gliomas, the most prevalent intracranial malignancies, account for approximately 81% of primary malignant brain tumors [1–3]. It typically appears on the cerebral hemispheres or brainstem, growing quickly and aggressively [1, 4]. Among them, glioblastoma multiforme (GBM), the highly heterogeneous and the most lethal high-grade glioma, with pronounced intra-tumoral heterogeneity and dismal prognosis: the median overall survival remains only 12 to 15 months [1, 3, 5–7]. Although standard treatment-maximal safe surgical resection followed by radiotherapy plus temozolomide chemotherapy-has modestly improved outcomes, the median survival has plateaued at 15-18 months, and the 5-year survival rate remains below 10% [2, 8, 9]. There is currently no cure largely due to the inherently heterogenous nature, therapeutic resistance, tumor recurrence, and the unique tumor microenvironment of the brain [3] , especially the dynamic interactions between glioma cells and neurons.

An upcoming concept has emerged in GBM biology: the recognition that the neural microenvironment and neuronal activity are not merely bystanders but active regulators of glioma initiation and progression [10, 11]. This concept is supported by several lines of evidence, including stemness signatures resembling neural development [12, 13], synaptic integration of glioma cells into neural circuits [14, 15] and the modulation of specific neurotransmitter or other neuronal secretory factors related pathways [16–18]. Several studies reported that artificially activated neuron in brain might provide the beneficial niche in promoting the initiation and development of glioma [10, 11, 19–22]. Physiological neuronal circuits in brain also contribute to the progression of glioma, including visual [22] and olfactory [21] pathway. All these findings establish a distinctive neuron-glioma cell microenvironment for GBM [23–25]. Conversely, glioma cells exert reciprocal influence on adjacent neurons, including secreting amounting glutamate causing neuronal excitotoxicity [26], perturbing chloride homeostasis in excitatory neurons and suppressing the function of gamma-aminobutyric acid (GABA) neurons [27–31]. All these alterations lead to excitation/inhibition imbalance in peritumoral networks, render the neuronal circuits hyperexcitable and prone to seizure activity [32]. Taken together, these bidirectional interactions underscore that clarifying how neuronal microenvironmental factors dictate glioma growth is an urgent frontier, and may uncover novel therapeutic vulnerabilities [23].

Emerging studies have begun to shed a new light on targeting neuronal activity to restrain glioma. For example, anti-seizure lamotrigine was found to block optic pathway glioma progression by modulating neuronal excitability [33]. Pharmacological inhibition of glutamatergic signaling prolongs survival in mice [3]. Given the new applications of anti-seizure medication and electrophysiological profile of glioma, we envisage that neuronal activity regulating medications might be repurposed as adjuvant anti-glioma therapies. Midazolam (MDZ), a benzodiazepine sedative, modulates GABA_A receptor-mediated inhibition and previous work has hinted the anti-cancer effects of MDZ in multiple tumors [34–36]. MDZ holds promise for disrupting aberrant neuron-glioma signaling.

We hypothesis that MDZ can suppress glioma growth by altering neuronal activity. In the present study, neuronal activity was modulated using potassium chloride (KCl) and MDZ, and the subsequent effects on glioma cell proliferation were investigated. The molecular mechanisms underlying this effect were further explored. Our findings confirm that MDZ-mediated suppression of neuronal activity attenuates glioma growth, suggesting a novel therapeutic approach targeting the neuron-glioma axis.

## Materials and Methods

### Primary cortical neurons isolation, culture and pharmacological treatments

Primary neurons were isolated from C57BL/6 E16-18 brains using a non-enzymatic mechanical dissociation method. Brains were moved into ice-cold PBS, and the cerebral cortex was dissected with meninges removed. Tissue was gently dissociated by repeated trituration using pipettes. The cell suspension was centrifuged at 1000 rpm for 5 min and resuspended in complete neuronal culture-medium consisting of Neurobasal medium, B27, Penicillin-Streptomycin (Thermo Fisher Scientific, USA) and L-Glutamine (Sigma, USA). Primary cortical neurons were coated on six-well plates at a density of 70,000 cells/cm^2^ for 5 days *in vitro* at 37 °C and 5% CO_2_. Neuronal depolarization was quantified as the percentage of c-Fos positive cells among MAP2 positive neurons (the percentage of c-Fos⁺ cell in MAP2⁺ cells). To collect depolarized neuronal conditional medium (Neu-CM), cortical neurons were treated with 10, 16 or 30 mM KCl (Macklin, China) for 4 h [37]. For MDZ (Jiangsu En Hua, China) treatments, cortical neuron were incubated with 30 mM KCl and followed by 5, 40 or 100 nM MDZ for 4 h, respectively [38]. After treatments, primary neurons were washed three times with PBS, and then incubated in fresh complete medium for an additional 24 h before Neu-CM were collected.

### Culture of glioma cells

GL261 and T98 glioma cells, obtained from Mou’s laboratory at Sun Yat-sen University Cancer Center, were cultured with medium containing high glucose Dulbecco’s modified Eagle’s medium (Hyclone, USA) and 10% fetal bovine serum (BOVOGEN, Australia) and 1% Penicillin-Streptomycin at 37 °C and 5% CO_2_. Glioma cells were washed twice with PBS and co-cultured with Neu-CM for 24 h for following experiments. All cell lines tested negative for contamination.

### 5-Ethynyl-2ʹ-Deoxyuridine (EdU) staining assay

The proliferation of glioma cells was determined by BeyoClick™ EdU Cell Proliferation Kit (Beyotime, China). Briefly, glioma cells were treated with Neu-CM for 24 h and EdU was added at 10uM for 2 h prior to cell harvesting. For glioma proliferation quantification, EdU^+^ cells were counted and normalized to the total number of DAPI^+^ nuclei cells (the percentage of EdU⁺ cell in DAPI⁺ cells) [19]. The EdU-labeled cells were fixed and then incubated with Click reaction mixture for 30 min at room temperature in the dark. The glioma cells were washed and stained with DAPI. The samples were analyzed for red fluorescence for EdU labeling and blue fluorescence for DAPI labeling with a fluorescence microscope (Leica Microscope DMi8 or Olympus IX73). Multiple fields of view were analyzed per sample, and experiments were performed with three independent biological replicates. Images were processed using ImageJ software. After 24 h incubation in Neu-CMs with or without 1ug/ml IGF1 binding protein-3 (Sigma, USA) [39] , 100 μM IGF 1R inhibitor Linsitinib (OSI-906) (Sellect, USA) [40, 41] or exogenous IGF1 factor (MCE, USA), the proliferation of glioma cells was detected by EdU assay.

### Real time-quantitative polymerase chain reaction (RT-qPCR)

RT-qPCR was performed as described previously [39]. Briefly, total RNA was isolated from cells using EZ-press RNA Purification Kit (EZBioscience, USA) according to the manufacturer’s instructions. and cDNA was generated by cDNA synthesis kit (TAKARA, Japan). RT-qPCR was utilized with TB green/Rox PCR Master Mix (TAKARA, Japan). All RT-qPCR reactions were carried out according to manufacturer’s instructions on Bio-Rad RT-qPCR system. Experiments were performed with three independent biological replicates. Detailed primer sequences for RT-qPCR are listed in Table 1.

**Table 1.**
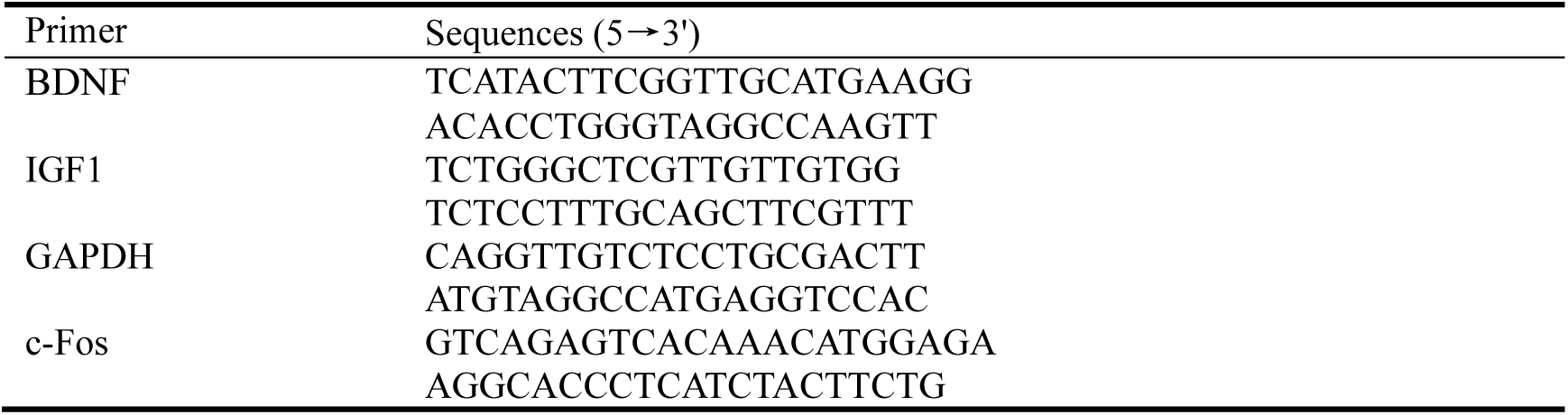
Sequences of primers used in the study.

### Western blotting

Glioma cells were collected and lysed in the cold RIPA buffer (Beyotime, China). Samples were centrifuged at 12,000g for 2 min at 4 °C and the supernatant was reserved. The concentration of total protein was normalized by the BCA protein quantification kit (Beyotime, China). Samples mixed with loading buffer were subjected to SDS-PAGE for electrophoresis, and the separated proteins were transferred to polyvinylidene fluoride membranes using the routine procedure. Polyvinylidene fluoride membranes were incubated with blocking buffer (Xblot, China) for 30 min and primary antibody solution at 4 °C overnight. After washing with TBST (3 times, 5 min each time), membranes were incubated with secondary antibodies coupled with horseradish peroxidase (diluted in TBST, peroxidase goat anti-mouse, 1:1,000; peroxidase goat anti-rabbit, 1:1,000.) for 1 Sh. Western blotting bands were detected by an ECL detection kit (Enzyme, China). The antibodies used in the study are detailed in Table 2.

**Table 2.** Antibodies used in the study.

| Target | Cat.No | Company |
| --- | --- | --- |
| IGFBP3 | SRP3067 | Sigma-Aldrich |
| PI3K | ET1608-70 | HUABIO |
| p-PI3K | HA721672 | HUABIO |
| AKT | 4691 | Cell Signaling Technology |
| pAKT (Ser473) | 4060 | Cell Signaling Technology |
| IGF1R | 9750 | Cell Signaling Technology |
| p-IGF1R | 3024 | Cell Signaling Technology |
| β-actine | ab8226 | Abcam |
| MAP2 | ab5392 | Abcam |
| NeuN | MAB377 | Sigma-Aldrich (Millipore) |
| c-Fos | 4384 | Cell Signaling Technology |
| Ki67 | ab15580 | Abcam |
| PDGFRα | ab203491 | Abcam |
| GFAP | N1506 | Dako |
| Donkey anti chicken-488 | 703-545-155 | Jackson ImmunoResearch Labs |
| Donkey anti rabbit-555 | A-31572 | Thermo Fisher Scientific, |
| Donkey anti mouse-488 | A-21202 | Thermo Fisher Scientific |
| Donkey anti Goat-555 | A-21432 | Thermo Fisher Scientific |
| HRP conjugated Goat Anti-Mouse IgG (H+L) | G1214 | Servicebio |
| HRP conjugated Goat Anti-Rabbit IgG (H+L) | GB23303 | Servicebio |

### Enzyme-linked immunosorbent assay (ELISA)

The concentrations of IGF1 in Neu-CMs were quantified using a commercial Mouse/Rat IGF1 ELISA Kit (Multi Sciences, China) according to the manufacturer’s instructions. Briefly, a total of 100 μL of standard or Neu-CMs sample was added to each well of a pre-coated microplate and incubated at room temperature for 2 h with shaking at 300 rpm. After washing six times, 100 μL of biotinylated detection antibody (1:100 dilution) was added and incubated for 1 h. Subsequently, the plate was washed and incubated with 100 μL of Horseradish Peroxidase (HRP)-conjugated streptavidin (1:100 dilution) for 45 min. After a final wash, 100 μL of TMB substrate was added and incubated for 30 min in the dark. The reaction was terminated with 100 μL of stop solution. The absorbance was measured at 450 nm (with a reference wavelength of 630 nm) using a microplate reader. The concentration of IGF1 was calculated based on a standard curve. All samples were measured in triplicate.

### Chromatin immunoprecipitation (ChIP) and ChIP-qPCR

ChIP assays were performed using primary neurons. Cells were cross-linked with 1% formaldehyde at room temperature for 10 min, quenched with 125 mM glycine for 5 min, and lysed in SDS lysis buffer containing protease inhibitors. Chromatin was fragmented by sonication using a Bioruptor system (Diagenode, Belgium) at medium power, with the water bath maintained at 2 °C, for 10 min. Chromatin shearing efficiency was verified by agarose gel electrophoresis. Equal amounts of chromatin were incubated overnight at 4 °C with anti-c-Fos antibody (CST, USA) or normal IgG as a negative control. Immune complexes were captured with Protein A/G magnetic beads, washed sequentially, eluted, and reverse cross-linked. ChIP DNA was purified using standard procedures. ChIP enrichment was quantified by RT-qPCR using SYBR Green chemistry. Experiments were performed with three independent biological replicates.

### Glioma xenografts model establishment and drug administration

The tumor xenografts were established in 6 to 8-week-old male C57BL/6 male mice obtained from Guangdong GemPharmatech Co., Ltd (China). All animal experiment protocols were approved by the Affiliated First Hospital of Guangzhou Medical University Institutional Animal Care and Use Committee (No.2022235). In a nutshell [42], mice were continuous anesthetized with 2% Isoflurane (RWD, China) and the skull of the mouse was exposed with a small opening using a 25 gauge needle (Gao Ge, China) 1.0 mm to the left of midline, 1.0 mm posterior to the bregma [43]. A total of 3× 10^5^ GL261-Luciferase-GFP glioma cells were injected into the brain (3.0 mm deep under the pia surface [43]). Glioma xenografted mice were randomized into two groups and intraperitoneally injected with PBS (vehicle control group, n = 7) or 16 mg/kg MDZ [34, 44] (MDZ-treatment group, n = 7) once every two days for 3 weeks. All mice were monitored every day for the development of symptoms related to tumor burden and body weight (g) were measured. Tumor growth was quantified using bioluminescence imaging with the IVIS Spectrum system. Regions of interest were defined over the tumor area, and bioluminescence signals were quantified as total photon flux (photons/s). For longitudinal analyses, bioluminescence signals were normalized to baseline values for each mouse. Mice were euthanized when they exhibited symptoms indicative of significant compromise to neurological function.

### Immunofluorescence

In brief [39], primary antibodies of cell and tissue immunofluorescence staining were listed in Table 2. Mice were anaesthetized (3 % Isoflurane) and perfused through the left cardiac-ventricle with cold PBS. Brains were then fixed with 4 % paraformaldehyde overnight at 4 ℃. Fixed tissues were cryoprotected in 30% sucrose and embedded into optimal cutting temperature prior to cryo-sectioning on a cryostat. Tissue slices were then rehydrated with PBS, blocked in blocking solution, containing 3% donkey serum in PBST (PBS + 0.3 % Triton X-100), for 1h and incubated with primary antibodies (diluted in blocking solution) overnight at 4℃. After 3 times washes, Alexa dye conjugated secondary antibodies were applied at 1:500 dilution rate for 1h at room temperature. Following 3 times washes with PBS and counterstaining with DAPI, tissue slices were mounted in 70 % glycerol with coverslips. Images were acquired on fluorescence microscope system (Leica Microscope DM6) and processed using ImageJ.

### RNA-Sequencing (RNA-seq)

Total RNA was isolated from primary neurons treated with KCl and MDZ, as well as from glioma and normal tissues treated with vehicle or MDZ, using TRIzol® Reagent (Invitrogen, USA). RNA integrity and purity were assessed by agarose gel electrophoresis, NanoPhotometer spectrophotometry, and an Agilent 2100 Bioanalyzer. RNA-sequencing was performed by Shanghai Origingene Bio-pharm Technology Co.Ltd and strictly obey standardized sequencing procedures. To minimize sequencing-related technical noise and redundancy, global sample consistency was evaluated using pairwise Pearson correlation across all features with correlation-based hierarchical clustering (distance = 1 - Pearson correlation) and principal component analysis (PCA) on the normalized expression matrix. Samples were retained for downstream analyses unless they showed both consistently reduced correlation with the remaining cohort and clear separation from the main sample cluster on the leading principal components. Differential gene expression analysis was performed using DESeq2 [45] with a *p*-value < 0.05 and |log_2_ fold change|> 0.7. Pathway enrichment analysis was performed using the cluster Profiler package with the Gene Ontology database. Pathways with an absolute normalized enrichment score (NES) greater than 1 and a false discovery rate (FDR) less than 0.05 were considered significantly enriched. Spearman correlation analysis was used to identify genes highly correlated with *Igf1* expression levels. To infer overall expression levels of genes sets, single sample gene set enrichment analysis (ssGSEA method from GSVA package) was used. Gene co-expression modules were identified through clustering analysis. For binary comparisons, partitioning around medoids (PAM) clustering was performed using the cluster package. For datasets with three or more groups, the Fuzzy c-means algorithm was implemented via the Mfuzz package [46]. The decoupleR [47] package univariate linear model was employed to infer IGF1 related transcription factor (TF) activity in each sample based on a curated collection of TFs and their transcriptional targets from the CollecTRI database and TFs were predicted by JASPAR database. All downstream bioinformatic analyses were performed by R version 4.2.2.

### Data and code availability

RNA-sequencing data of cortical neurons were deposited in GEO: GSE228253, and RNA-sequencing data of tumor and adjacent normal brain tissues in https://doi.org/10.6084/m9.figshare.30397444.v1. This study did not generate any unique code.

### Statistical Analyses

All the data are presented as the mean ± SEM. Normality of the data was assessed using the Shapiro-Wilk test. Data that passed the normality test were analyzed using parametric tests (Student’s t-test or one-way ANOVA). Unpaired Student’s t tests were performed to compare the means of the two groups. One-way ANOVA was used for comparisons among the different groups that involved one factors. Statistical analyses were performed via GraphPad Prism 6.0. *P* value of <0.05 was considered statistically significant.

## Results

### Activated neuronal activity promotes glioma cells proliferation

To investigate whether neuronal activation promotes glioma progression, primary cortical neurons were depolarized for 4 h with increasing concentrations of KCl (0, 10, 16, and 30 mM) to activate neuronal activity, and the Neu-CMs were subsequently collected for co-culture with glioma cells (Figure 1A). Microtubule-Associated Protein 2 (MAP2) is used as a marker of mature neurons, whereas c-Fos serves as an indicator of neuron activation. The proportions of activated neurons in these groups were approximately 0.4%, 8%, 39%, and 78%, respectively, indicating that 30 mM KCl elicited robust neuronal activation (Figure 1B-D). When glioma cells were cultured with depolarized Neu-CMs, a marked increase in proliferation was observed, as determined by EdU incorporation assays. The proliferation rates of both GL261 and T98 glioma cell lines positively correlated with the degree of neuronal activation, suggesting that depolarized Neu-CMs promote glioma cells entry into the S phase of the cell cycle (Figure 1E–G). In addition, the EdU fluorescence intensity exhibited dose-depended effect associated with neuron activation status, with 30mM KCl-exposed glioma cells showing the strongest proliferative effect. These results indicate that depolarized Neu-CM promotes glioma cells proliferation by accelerating DNA synthesis. To further ask whether this phenomenon is specific to glioma cells, PC9 (non-small cell lung cancer, NSCLC) cells were used and there was no obvious cell proliferation effect in PC9 under identical conditions (Supplementary Figure S1A-B). Collectively, these findings demonstrate that activated neuronal activity facilitates glioma cells proliferation and may serve as a potential therapeutic target for glioma intervention.

**Figure 1.**
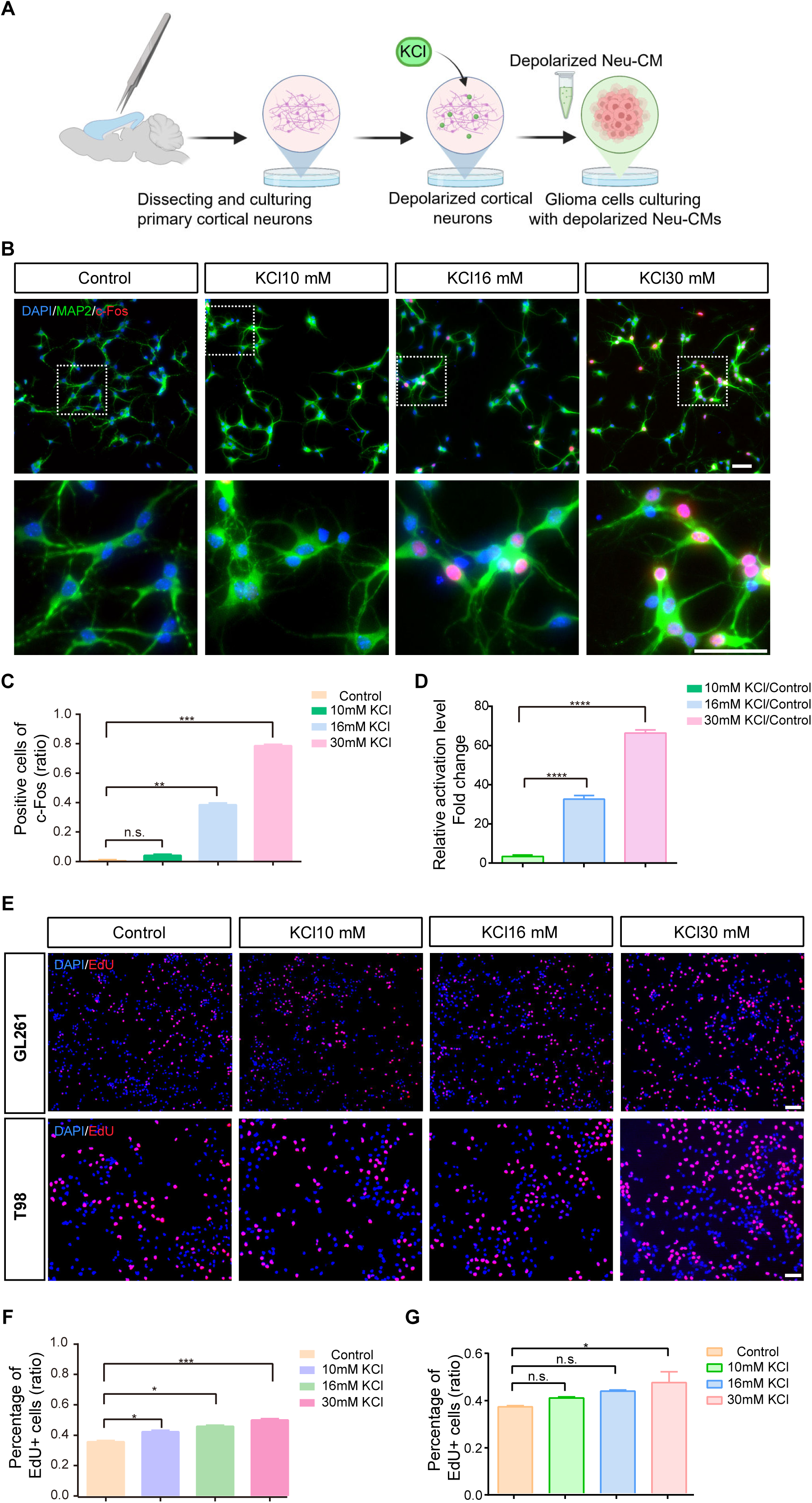
Depolarized cortical neurons promotes the proliferation of glioma cells. **(A)** Schematic illustration of depolarized neuron-glioma cell co-culture system. **(B)** Cortical neurons activated by four different concentrations of KCl (Control, 10, 16 or 30mM KCl). Scale bar = 50 µm. **(C-D)** Quantification of absolute and relative expression of c-Fos and MAP2 in depolarized neurons. \*\**P*< 0.01, \*\*\**P*< 0.001, \*\*\*\**P*< 0.0001, n.s *P*>0.05. n=3. **(E-G)** Cell proliferation of GL261 and T98 glioma cells co-cultured with depolarized neuronal conditioned medium were assessed via EdU assay. Scale bar = 100 µm. \**P*< 0.05, \*\*\**P*< 0.001, n.s. *P*>0.05. n=3.

### Midazolam attenuates glioma growth by suppressing neuronal activity *in vitro*

Next, to determine whether inhibition of neuronal activity could attenuate glioma proliferation, we employed MDZ to modulate neuronal excitability. Primary cortical neurons were treated with different concentrations of MDZ (5, 40, and 100 nM) to evaluate its inhibitory effect on neuronal activation (Figure 2A). The proportion of c-Fos^+^ neurons decreased in a concentration-dependent manner (0.42, 0.38, and 0.08, respectively), with 100 nM MDZ showing the strongest inhibitory effect (Figure 2B-D). Moreover, these MDZ concentrations did not noticeably alter synaptic morphology compared with the negative control or 30 mM KCl-treated groups. These results indicate that 100nM MDZ effectively suppresses neuronal activity with minimal morphological disruption, whereas higher concentration (500 nM) induces cortical neuron death (Supplementary Figure S2). Subsequently, MDZ Neu-CMs were used to co-culture with glioma cells. Just as we predicted, the proliferation of both GL261 and T98 glioma cells were significantly inhibited (Figure 2E-G). Glioma cell proliferation was inversely correlated with neuronal activity, suggesting that MDZ attenuates glioma growth by suppressing neuronal excitation and diminishing neuron-derived pro-tumor factors. Taken together, these findings demonstrate that pharmacological suppression of neuronal activity with MDZ effectively reverses the pro-tumor effect of hyperactive neurons, highlighting the potential of neuronal activity modulation as a therapeutic approach for glioma.

**Figure 2.**
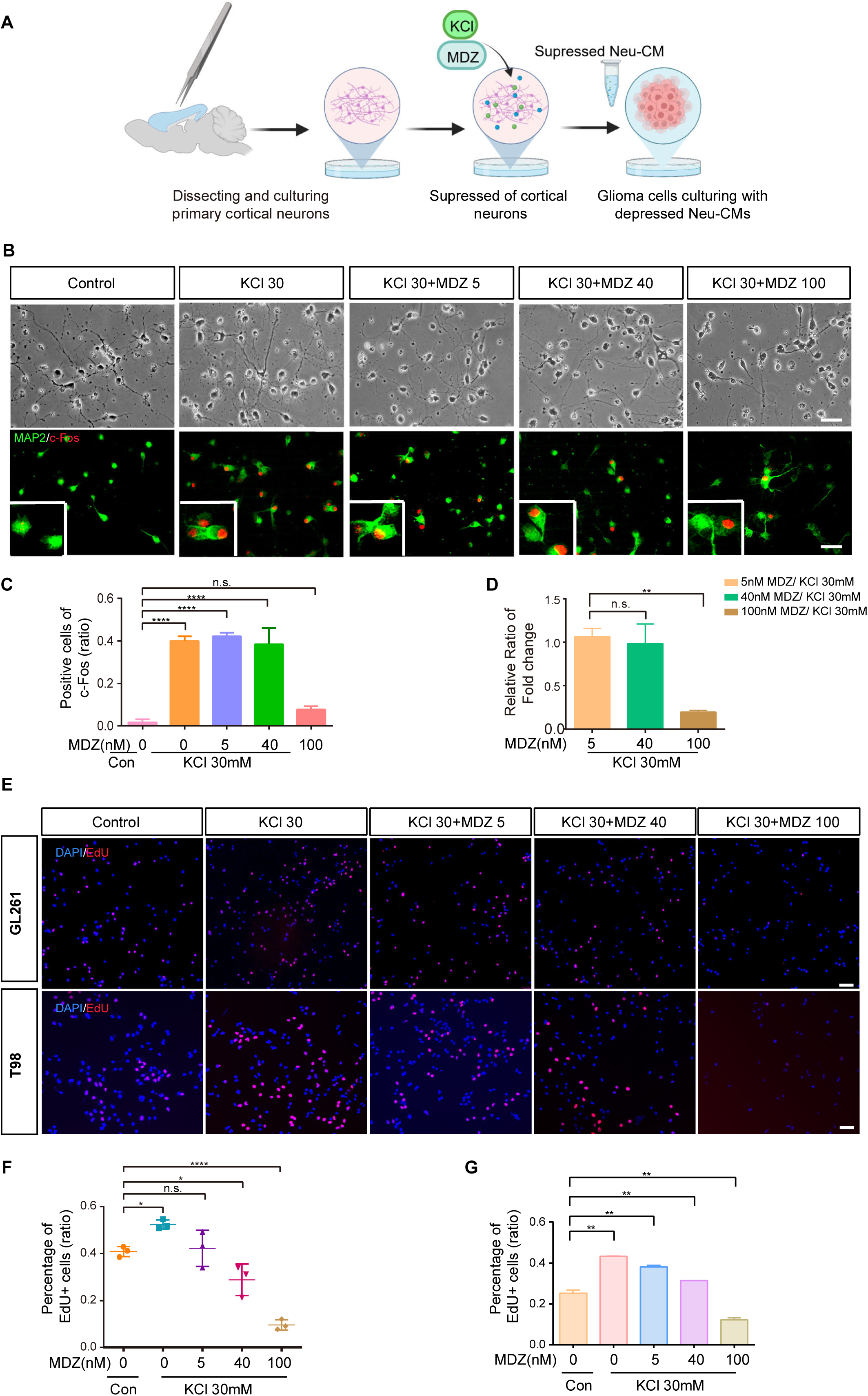
Suppressed cortical neurons inhibit the proliferation of glioma cells. **(A)** Schematic illustration of suppressed neuron-glioma cell co-culture system by MDZ. **(B)** Representative images showing morphology and activation level changes of cortical neurons treated with 30 mM KCl plus 5, 40 or 100nM MDZ. Scale bar = 100 μm. **(C-D)** Quantification of absolute and relative levels of neuronal activation after MDZ treatment, as assessed by c-Fos immunofluorescence. \*\*\*\**P<* 0.0001, n.s. p>0.05, n=3. Representative EdU staining **(E)** and quantification **(F-G)** showing proliferation of GL261 and T98 glioma cells co-cultured with CM from MDZ-treated neurons. Scale bar = 100 µm. n.s. *P*>0.05, \**P*< 0.05, \*\**P*< 0.01, \*\*\*\**P*< 0.0001. n=3.

### Neuronal activity-dependent expression of IGF1 is essential for glioma cell proliferation

To investigate the underlying mechanism of MDZ-mediated inhibition of glioma cells proliferation, RNA-seq was performed on cortical neurons under depolarized or suppressed conditions, subsequently accompanied by bioinformatic analyses (Figure 3A). The expression of brain-derived neurotrophic factor (BDNF) [48], the first recognized neuronal activity-dependent factor, was regarded as a canonical gene marker for validating the effectiveness of KCl and MDZ in modulating neuronal activity (Figure 3A-B). A similar pattern was observed for IGF1, indicating that its expression is regulated by neuronal activity (Figure 3A-B). To validate the sequencing results, quantitative RT-qPCR was conducted to assess the expression of IGF1 and BDNF. Both of them were increased in the depolarized group and decreased in the MDZ-suppressed group (Figure 3C). IGF1 is a well-characterized growth factor that stimulates mitosis, inhibits apoptosis, and regulates cellular metabolism [49, 50]. To determine whether neuron-derived IGF1 is essential for glioma proliferation, IGF binding protein 3 (IGFBP3) was added to neutralize IGF1 activity in Neu-CMs. The proliferative capacity of co-cultured glioma cells was dramatically reduced following IGF1 blockade (Figure 3D-E). Consistent with the findings from IGFBP3 results, pharmacological blockade of the IGF1R with Linsitinib (OSI-906) significantly abolished the proliferative effect on glioma cells (Figure 3F-G). The inhibitory efficacy of Linsitinib was validated by the downregulated phosphorylation of the IGF1R signaling pathway, as evidenced by the decreased ratio of p-IGF1R/IGF1R (Supplementary Figure S3A-B). Furthermore, we confirmed that IGF1 secretion was elevated in the Neu-CM of depolarized neurons (Supplementary Figure S3C). Correspondingly, the addition of exogenous recombinant IGF1 protein directly promoted glioma cell proliferation, as indicated by a significant increase in the percentage of EdU^+^ cells (Supplementary Figure S3D-E). Consistently, the phosphorylation level of PI3K and AKT, key downstream mediators of the IGF1 signaling pathway [51, 52], were increased in glioma cells co-cultured with depolarized Neu-CM and decreased in those cultured with suppressed Neu-CM (Figure 3H-I). In addition, PI3K/AKT signaling was also decreased in the IGFBP3 experiment (Figure 3J-K). Collectively, these results demonstrate that neuronal activity-dependent IGF1 plays an essential role in promoting glioma cell proliferation [21] via activation of the PI3K/AKT pathway, consistent with previous findings [53].

**Figure 3.**
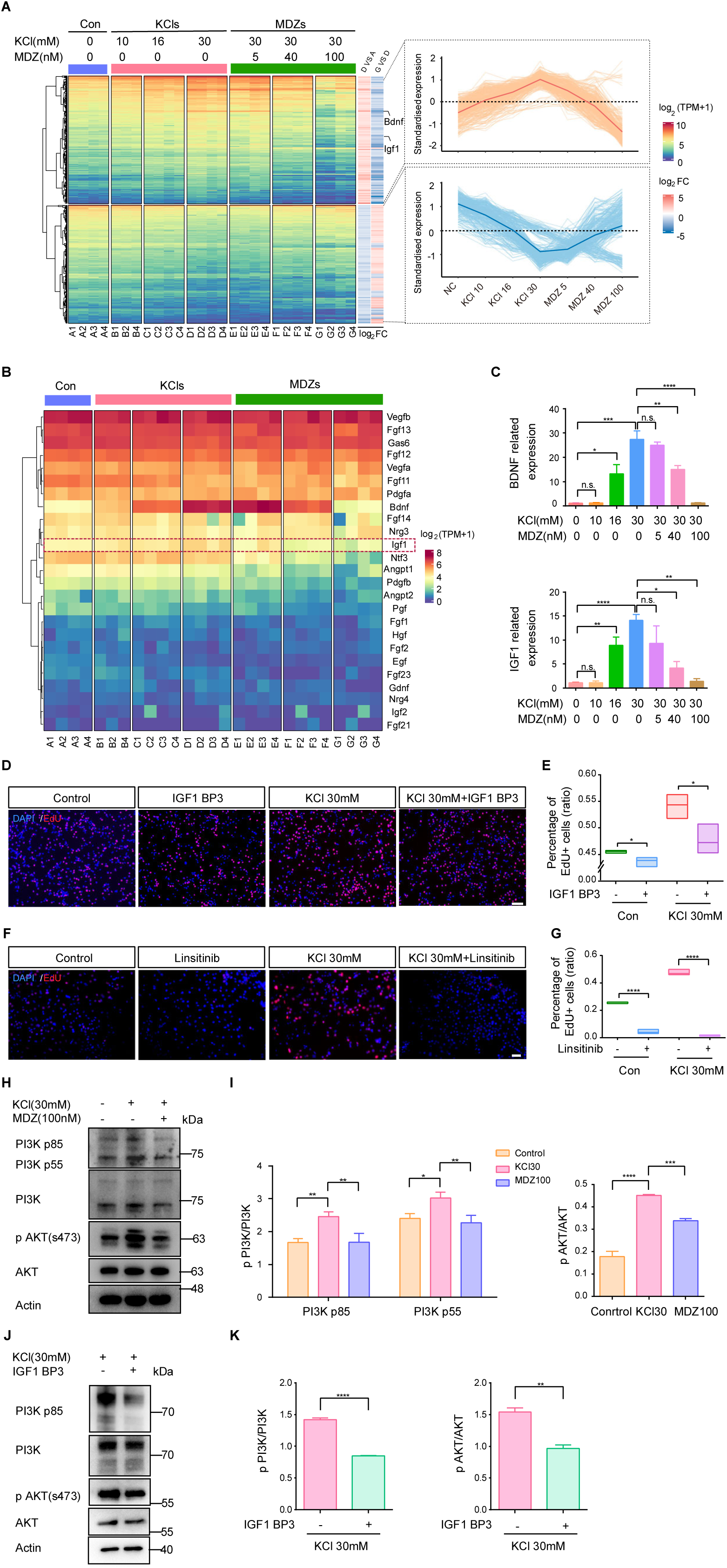
IGF1 functions as an essential neuronal activity-dependent growth factor promoting glioma cells proliferation. **(A)** Differentially expressed genes and overall expression trends of cortical neurons from seven different groups. “D vs A”: log₂FC of KCls vs Control, “G vs D”: log₂FC of MDZs vs KCls. **(B)** Gene expression profiles of commonly secreted neuronal factors under KCl, and MDZ conditions. IGF1 is highlighted by a red dashed line. **(C)** Validation of IGF1and BDNF expression by RT-qPCR. n.s. *P*>0.05, \**P*< 0.05, \*\**P*< 0.01, \*\*\*\**P*< 0.0001. n=3. **(D-E)** Representative images and quantification of EdU assay after IGFBP3 experiment in GL261 cells. Scale bar = 100 μm. **(F-G)** Representative images and quantification of EdU assay after Linsitinib (OSI-906) treated experiments in GL261 cells. Scale bar = 200 μm. **(H-I)** Activation of the IGF1 downstream PI3K/AKT signaling pathway in glioma cells and its quantitative analysis. **(J-K)** PI3K/AKT signaling pathway change after IGFBP3 incubation in glioma cells. n.s. *P*>0.05, \**P<*0.05, \*\**P<* 0.01, \*\*\**P<* 0.001, \*\*\*\**P<* 0.0001. n=3.

### Neuronal activity regulates *Igf1* transcription through the transcription factor c-Fos

Since we found the change of IGF1 expression in a neuronal-activity-dependent way, we next want to confirm that IGF1is transcriptionally regulated by neuronal activity. To elucidate the signal transduction mechanisms underlying IGF1 regulation, we identified genes that co-express with *Igf1* expression (Figure 4A). Functional enrichment analysis revealed that IGF1-correlated genes were predominantly involved in neuroactive ligand-receptor interaction, axon guidance, and MAPK signaling pathway, whereas negatively correlated genes were enriched in necroptosis and carbon metabolism (Figure 4B). Given the distinct association patterns, we next examined whether these pathways were regulated by neuronal activity. Consistent with this possibility, KEGG pathway analysis revealed similar activity-dependent trends. Pathways related to neuroactive ligand-receptor interaction and MAPK signaling were upregulated in the KCl-treated (depolarized) group (Figure 4C, Supplementary Figure S4) and suppressed following MDZ treatment (Figure 4D, Supplementary Figure S4). Notably, MAPK signaling is a well-established pathway involved in neuronal activation and activity-dependent gene regulation [54]. To further explore activity-associated transcriptional programs, we performed clustering analysis of genes within the MAPK signaling pathway. This analysis identified five distinct expression patterns, among which genes in cluster 2 exhibited expression changes most closely aligned with neuronal activity modulation (Figure 4E). Given that immediate early genes such as *Fos* are classical downstream readouts of activity-dependent MAPK signaling, these results prompted us to further investigate the involvement of c-Fos in the regulation of *Igf1* transcription. Further analysis using a univariate linear model was performed to infer IGF1-associated transcription factor activity in each sample. *Igf1* expression showed a strong association with both the expression level and inferred transcriptional activity of Fos (Figure 4E-F). Analysis using the JASPAR database identified putative c-Fos binding motifs within the *Igf1* promoter region, suggesting a potential transcriptional association between c-Fos and IGF1 (Figure 4G). This prediction was further supported by ChIP-qPCR analysis, which confirmed c-Fos occupancy at the *Igf1* promoter (Figure 4H). Collectively, these results indicate that neurons respond to environmental stimulation by engaging activity-dependent transcriptional programs, and that c-Fos acts as a key transcriptional mediator linking neuronal activation to *Igf1* transcription.

**Figure 4.**
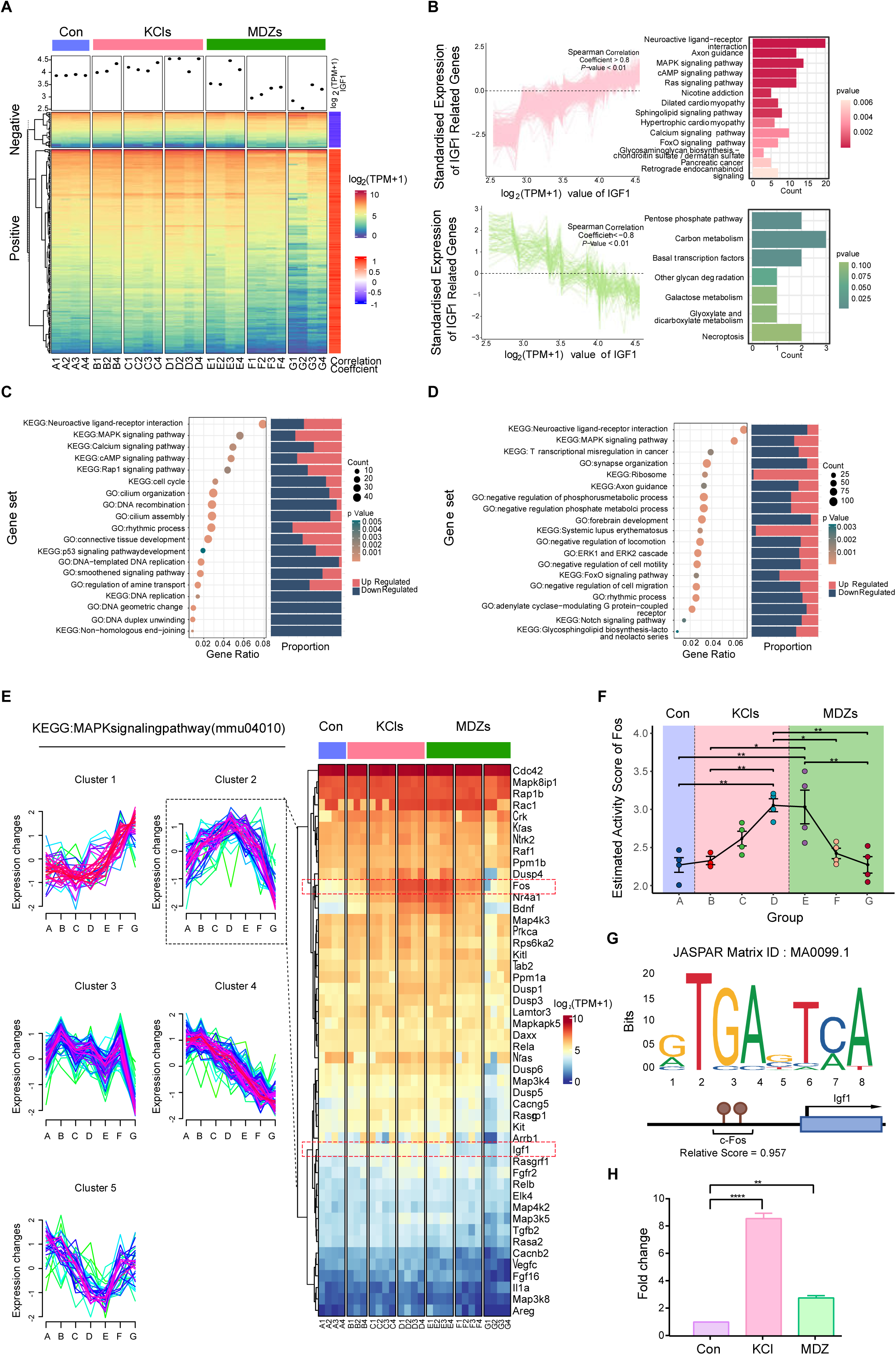
IGF1 is closely associated with neuronal activity. **(A)** Overall distribution of IGF1*-*related genes in all samples. **(B)** Standardized expression profile and function annotation of IGF1co-expression genes. **(C-D)** Over-representation enrichment analysis of genes of interest using Gene Ontology (GO) and Kyoto Encyclopedia of Genes and Genomes (KEGG) databases in KCl treated group and MDZ treated group, respectively. **(E)** Fuzzy cluster analysis identifying five distinct expression patterns of genes participated in MAPK signaling pathway. **(F)** Transcriptional activity of Fos was evaluated using a univariable linear model. **(G)** Predicted c-Fos binding motif within the *Igf1* promoter region identified using the JASPAR database. **(H)** ChIP-qPCR analysis assessing c-Fos enrichment at the predicted regulatory region. \**P*<0.05, \*\**P*< 0.01, \*\*\*\**P<* 0.0001. n=3.

### Midazolam suppresses glioma progression *in vivo*

Next, we investigated whether MDZ could inhibit glioma *in vivo* by suppressing neuronal activity. Glioma xenografted mice were intraperitoneally administered MDZ or vehicle respectively (Figure 5A). The sedative effect of MDZ was first confirmed by behavioral observation, and a supplementary video was recorded to document the activity changes (Supplementary Video). MDZ-treated mice entered in a quiescent state, whereas vehicle-treated mice remained freely active. Suppression of neuronal activity by MDZ was further validated by c-Fos immunofluorescence staining. Not surprisingly, the percentage of activated neurons (c-Fos^+^ cells/ NeuN^+^ cells) was significantly decreased in the MDZ group compared with the vehicle group (Figure 5B-C). Bioluminescence imaging using the IVIS Spectrum system demonstrated that MDZ treatment markedly reduced tumor growth relative to vehicle control, indicating a clear inhibitory effect on glioma progression (Figure 5D-E). Consistently, MDZ-treated mice exhibited better overall physical condition and higher activity levels than those in the vehicle group. During the experimental period, MDZ-treated mice body weight increases obviously were comparable to those of vehicle-treated controls (Figure 5F). As we expected, both the tumor volume (Figure 5G-H) and the proliferative index (Ki67^+^ cells/DAPI^+^ cells) (Figure 5I-J) were significantly decreased following MDZ administration. Some research studies have found that astrocytes become reactive, changing their function and supporting the tumor invasion [55, 56]. In our study, astrocytes were also found around the tumor to restrict glioma growth in MDZ group, supporting the notion that MDZ attenuated glioma growth by regulating tumor microenvironment (Supplementary Figure S5). Our results indicate that MDZ suppresses tumor microenvironment neuronal activity and consequently inhibits glioma progression *in vivo*, highlighting the therapeutic potential of targeting neuronal activity in glioma treatment.

**Figure 5.**
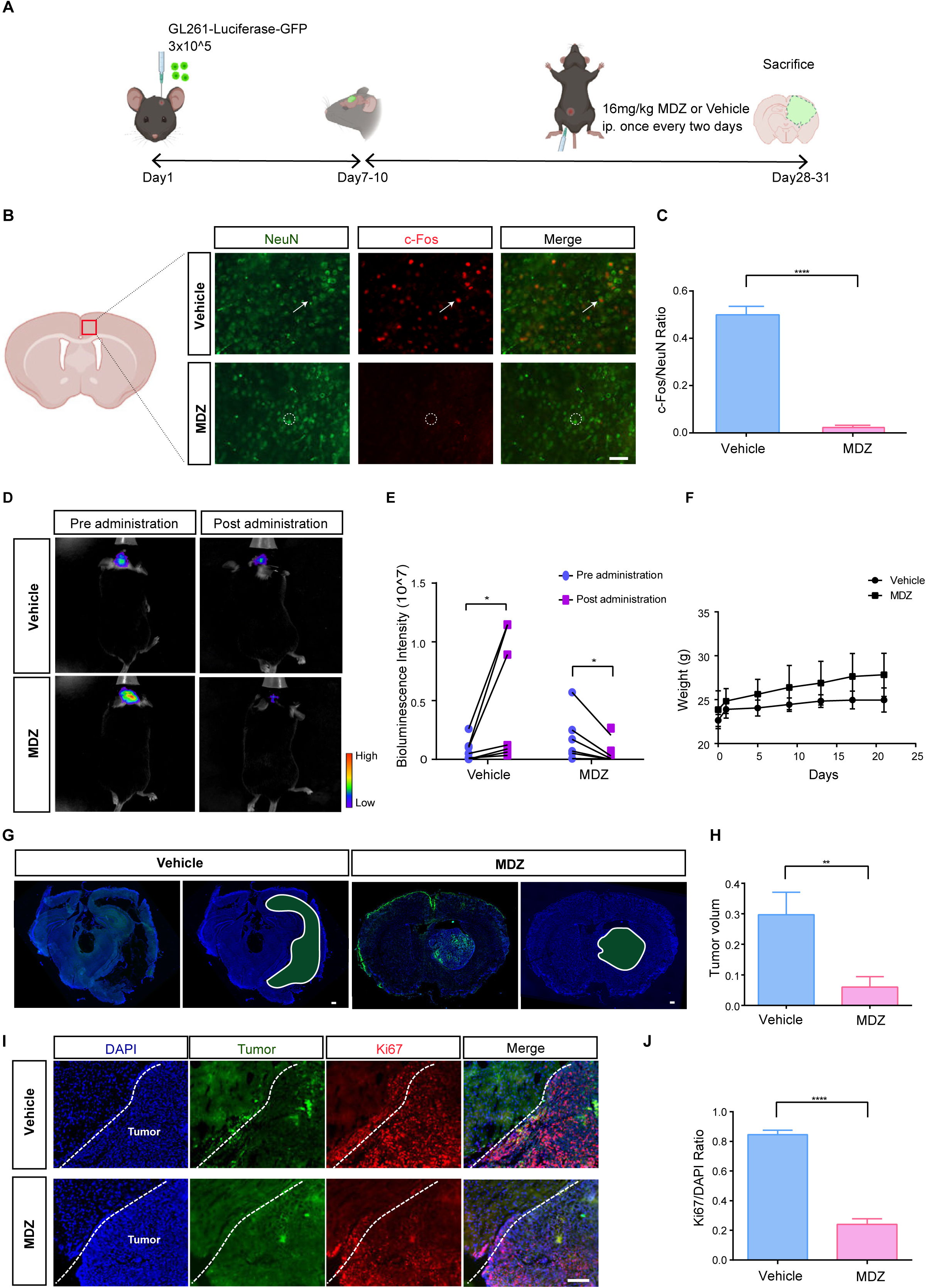
MDZ suppresses the neuronal activity and inhibited tumor growth *in vivo*. **(A)** Experiments design of glioma xenografted mouse model and treatment protocol. **(B-C)** Cortical neurons activity in mice treated by 16 mg/kg MDZ. Arrows point to the activated neurons and dotted circles denote suppressed neurons. Scale bar = 50 μm. **(D-E)** *In vivo* bioluminescence imaging and quantification of intensity before and after vehicle (ip. saline) or MDZ administration (n = 7). **(F)** Weight changes of mice in each group (n = 7). **(G-H)** Representative images and quantification (tumor area/ brain area) of tumor volume. White outlines and green fillings depict glioma boundaries. Scale bar = 300 μm. **(I-J)** Immunofluorescence staining and quantification of Ki-67 expression. Tumor regions are located to the right of the dotted line. Scale bar = 100 μm. * *P* < 0.05, \*\**P* < 0.01, \*\*\*\**P*< 0.0001. n=3.

MDZ globally suppresses neuronal activity-related transcriptional programs in glioma tissue To further elucidate the molecular basis underlying the inhibitory effect of MDZ on neuronal activity and glioma progression, transcriptomic profiling was performed on tumor and adjacent normal brain tissues from MDZ-treated and vehicle-treated mice. Heatmap analysis demonstrated distinct gene expression patterns between the MDZ-treated and vehicle groups. In adjacent normal tissues, MDZ markedly suppressed the expression of genes associated with activity (*Fosl2, Fosb, Fos*), neurotrophic growth factors (*Bdnf, Tgfb3, Ngf*), neuron-tumor supportive factors (*Snap25, Slit2, Efnb2*) and excitatory neuronal channels (*Gria2, Grik4, Grik5*) (Figure 6A).A similar trend was observed in tumor tissues, MDZ markedly suppressed the activity related and neuron-tumor supportive genes (Figure 6B), suggesting a shift toward reduced neuronal excitability in MDZ treated group. Consistently, pathway enrichment scores highlighted decreased activity of synaptic vesicle fusion, neurotransmitter receptor regulation, and glutamate signaling pathways, together with upregulation of immune antigen-processing signatures in the MDZ group (Figure 6C). In addition, precursor proliferation and synaptic transmission were significantly downregulated following MDZ treatment compared with the vehicle group (Figure 6D). These results indicate that MDZ reshapes both immune and neuronal transcriptional landscapes within the tumor microenvironment. Given that our central hypothesis concerns the modulation of neuronal activity by MDZ and its impact on glioma progression, we therefore validated the RNA-seq findings by examining neuronal activity signals in both tumor and adjacent normal brain tissues. Specifically, we assessed c-Fos expression as a well-established marker of neuronal activation [54]. The reduction of c-Fos^+^ cells was obviously in MDZ treatment group, no matter whether in adjacent normal brain tissue or in tumor tissue (Figure 6E-F), consistent with our earlier observations (Figure. 5B). Collectively, these results indicate that systemic MDZ treatment is associated with reduced neuronal activation and decreased expression of neurotrophic signaling both within and surrounding glioma lesions, thereby contributing to its overall anti-tumor effect.

**Figure 6.**
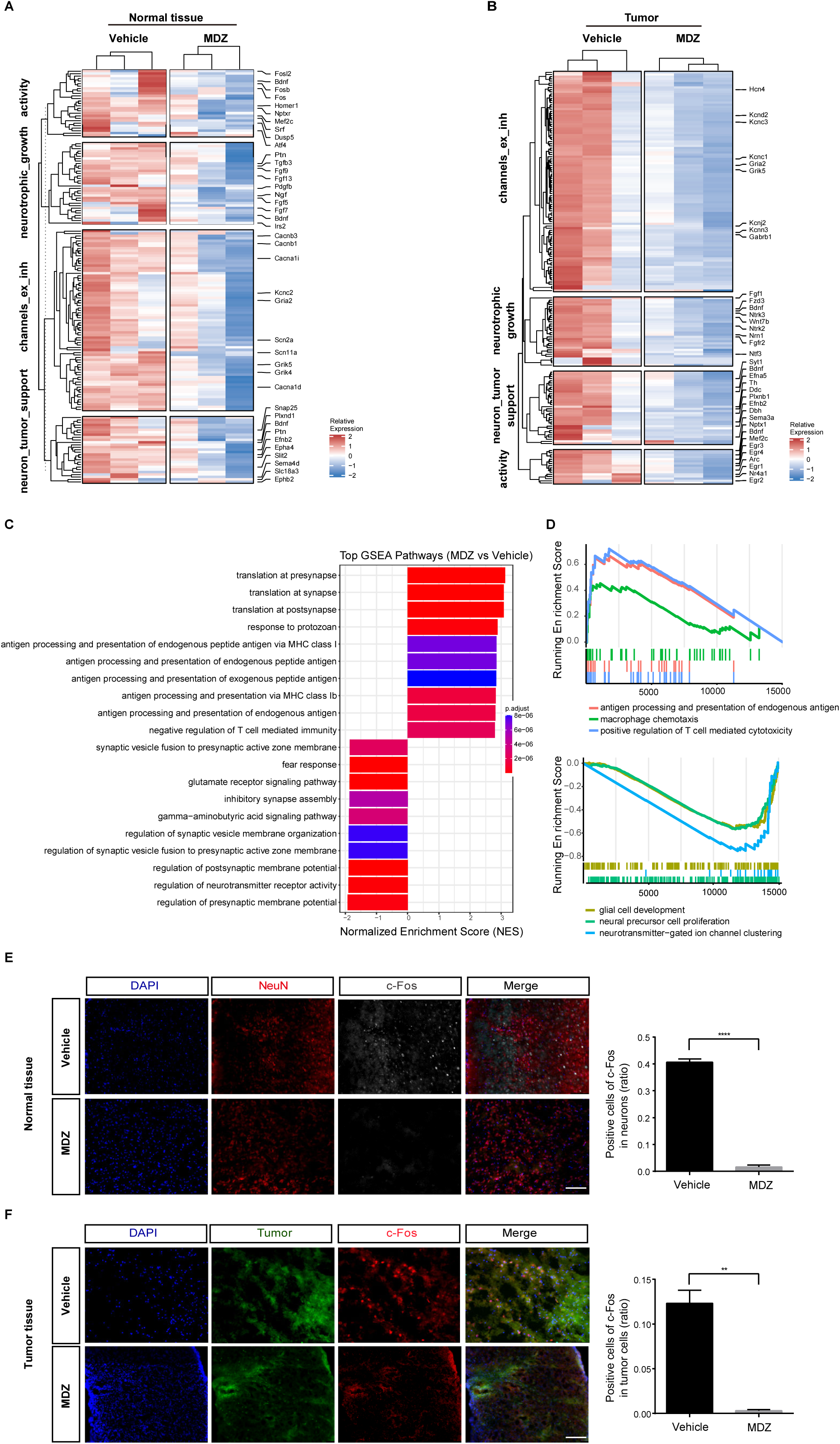
IGF1 is the essential pro-tumor growth factor for neuron-glioma co-culture system. **(A)** Heatmap of adjacent normal brain tissue showing transcriptional changes between vehicle- and MDZ-treated groups. **(B)** Heatmap of glioma tumor tissue showing transcriptional changes between the two groups. **(C)** T Top enriched pathways identified by GSEA in tumor tissue. **(D)** GSEA showing immune process and neuronal-activity-associated pathways. **(E-F)** Representative images and quantification of c-Fos in adjacent normal tissue and tumor tissue, respectively. Scale bar = 100 μm. \*\**P* < 0.01, \*\*\*\**P*< 0.0001.

## Discussion

The exploration of neuroscientific directives in oncological interventions [14, 15, 20, 57], while promising, presents a myriad of complexities and challenges, as neuronal signaling is essential for normal brain homeostasis [10]. However, targeting neuronal activity for therapeutic purposes remains challenging. In the present study, MDZ demonstrates a potential inhibitory effect on glioma progression by modulating neuronal activity both *in vitro* and *in vivo* experiments (Figure. 7).

**Figure 7.**
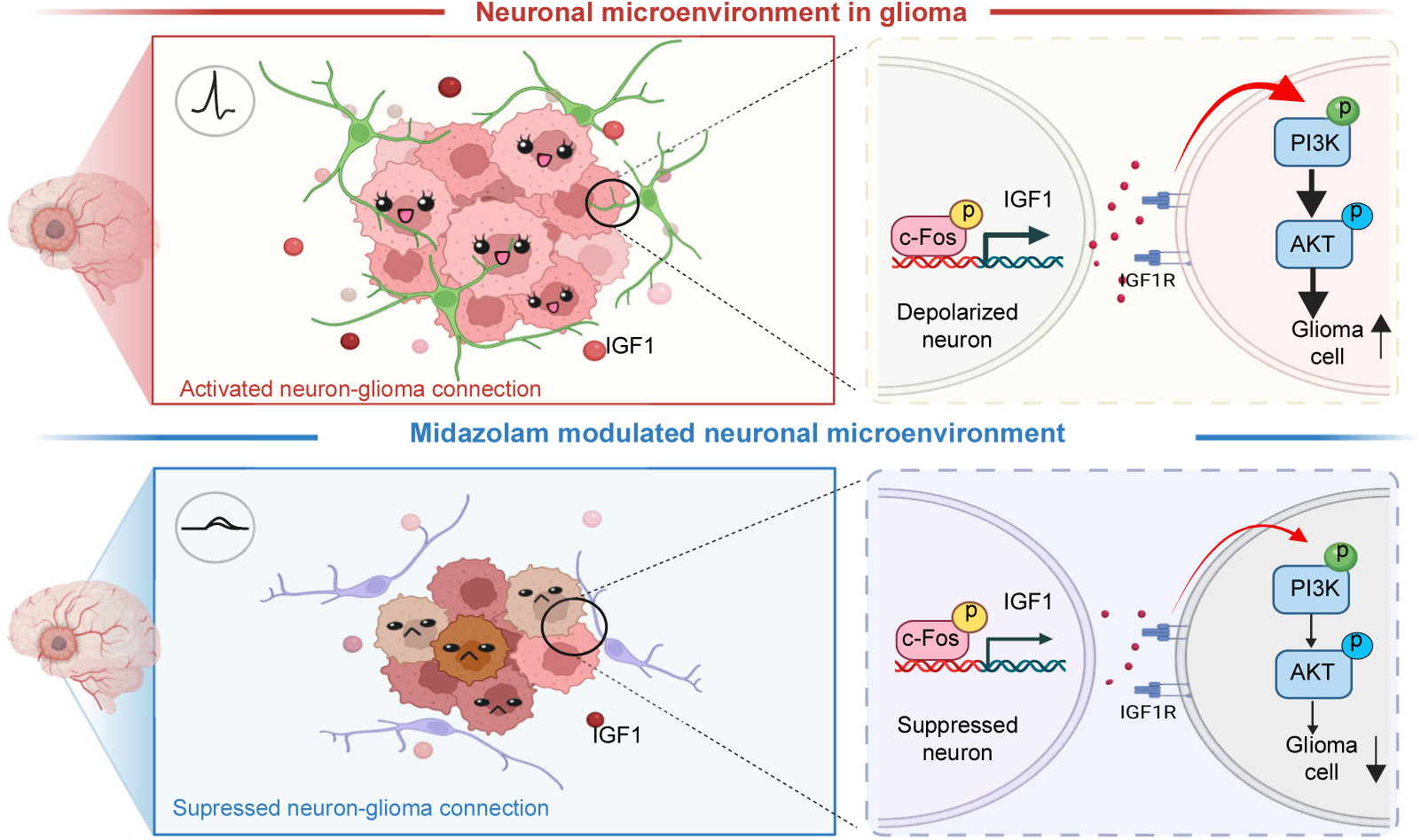
Proposed model for the inhibitory effect of MDZ on glioma progression. MDZ suppresses glioma progression by attenuating neuronal activity and downregulating IGF1secretion. This reduction dampens the activation of the PI3K/Akt signaling cascade in glioma cells, thereby impeding their proliferative capacity.

Recent evidence highlights the pivotal role of GABAergic signaling in glioma biology [30]. Interestingly, recent studies have reported that benzodiazepines may accelerate the proliferation of H3K27M^+^ diffuse midline gliomas (DMGs) by enhancing GABAergic currents [29]. This discrepancy likely reflects cellular context: This because DMG cells express high levels of the Na-K-2Cl cotransporter [29], leading to an efflux of Cl^-^ upon GABAARs and resulting in depolarization rather than inhibition[29]. IGF1 has been identified as an indispensable factor to promote glioma-genesis [21, 58]. Genetic or pharmacological interference with IGF1R activity inhibits the growth and progression of glioma cells in preclinical mouse models [21, 58], further supporting the therapeutic relevance of IGF1/PI3K/AKT signaling. Our findings demonstrate that MDZ reduces IGF1 expression in a neuronal-activity-dependent manner, suggesting that sedative modulation of neuronal excitability may indirectly restrain glioma growth via the IGF1 axis.

Growing evidence suggests that GBM secretes excessive excitatory neurotransmitters, such as glutamate, leading to neuronal hyperexcitability, epileptic activity, and neuropsychiatric symptoms [59]. This heightened neuronal activity not only promotes tumorigenesis but is also implicated in the onset of epileptic episodes and other neuropsychiatric manifestations [60–63]. MDZ has been documented for its efficacy in mitigating seizure activity and addressing sleep and anxiety disorders [64, 65]. Such findings underscore the potential applicability of MDZ in the management of both central and peripheral solid tumors. Notably, MDZ has been shown to modulate the miR-194-5p/HOOK3 signaling pathway, thereby potentiating cisplatin sensitivity in cisplatin-resistant NSCLC cases [66]. Its dual properties-attenuating neuronal activity and mitigating neuropsychiatric symptoms-make it a compelling candidate for adjuvant therapy in glioma patients. Importantly, given the efficient penetration across the blood-brain barrier of MDZ, it holds promise as a vehicular agent or as a foundation for derivatized substrates aimed at facilitating targeted drug delivery within the central nervous system. Collectively, these findings suggest that MDZ may provide multifaceted therapeutic benefits, offering both neuroprotective and anti-tumor effects in glioma management.

Even though gliomas can be inhibited by blocking neurotrophic factors and neurotransmitters, the neuronal network excitation remains extremely high, consistently providing a supportive niche for glioma cells. Even if the neurotrophic “seeds” are temporarily eliminated, the “soil” for tumor regrowth remains fertile. Therefore, more comprehensive management and new treatment strategy are urgently needed to deal with this disaster disease. Notably, recent translational studies highlight the promise of neuro-modulatory approaches in glioma therapy. For instance, the FDA-approved antidepressant treats incurable brain cancer in preclinical trial [2], and the neuroligin 3-release inhibitor INCB7839 is currently in a phase I clinical trial for children with recurrent or progressive high-grade glioma. These advances underscore the great potential of targeting neuronal activity and neuronal factor as a new direction for glioma treatment.

While this study demonstrates that MDZ suppresses glioma progression by attenuating neuronal activity and downregulating IGF1 signaling, several limitations should be acknowledged. First, most mechanistic evidence is derived from mouse models and *in vitro* experimental systems, and therefore a translational gap between animal models and human glioma physiology remains. Second, although multiple glioma models were examined, gliomas are highly heterogeneous tumors. Whether the neuron-IGF1-glioma axis identified here is broadly applicable across distinct glioma subtypes or molecular backgrounds remains to be determined. Third, the present study primarily focused on short-term modulation of neuronal activity and IGF1 signaling. Potential long-term adaptive responses, compensatory mechanisms, or the development of tolerance or resistance following prolonged MDZ exposure were not addressed and warrant future investigation.

Beyond its established sedative and neuroprotective properties, MDZ may represent a promising therapeutic adjuvant for GBM, contributing not only to tumor control but also to the alleviation of neurological symptoms. As the field of oncology evolves, the emphasis is increasingly shifting towards improving the quality of life of patients, rather than solely focusing on tumor eradication. Therefore, MDZ alone is unlikely to achieve curative efficacy, its integration into multimodal therapeutic strategies may improve both survival and quality of life for glioma patients, reflecting a shift toward more holistic approaches in neuro-oncology.

## Acknowledgments

This study was supported by grants from the National Natural Science Foundation of China (Grant Nos.82272224, 32470888), the Basic and Applied Basic Research Foundation of Guangdong Province (Grant Nos. 2021A1515220042), Natural Science Foundation of Guangdong Province (Grant Nos. 2022A1515012475), Guangzhou Science and Technology Plan Project -2024 Joint Funding Project by Municipal Universities (Research Institutes) and Enterprises (Grant Nos. 2024A03J1221), Guangzhou Science and Technology Program, 2024 Basic and Applied Basic Research Project (Young Doctor “Qihang” Direction) (Grant Nos.2024A04J3319).

---

**Supplementary Figure S1.** Representative imaging and quantification of EdU incorporation in PC9 cells under different medium conditions. The Roswell Park Memorial Institute (1640) medium served as medium control group, the left four groups were based on the Neu-CMs containing the concentration of KCl 0,10,16 and 30 mM, respectively. \*\**p*≤ 0.01. n = 3

**Supplementary Figure S2.** Representative imaging of 500 nm MDZ induced cortical neuron morphological disruption. Scale bar = 100 μm.

**Supplementary Figure S3.** (A-B) p-IGF1R/IGF1R signaling change in Linsitinib (OSI-906)-treated glioma cells. \*\*\*\**P<* 0.0001. n=3. **(C)** Relative results of ELISA analysis of IGF1 concentration in depolarized Neu-CM. n.s. \**P<* 0.05. n = 3. **(D)** Representative imaging and quantification of EdU incorporation in GL261 cells after the addition of recombinant IGF1. \*\*\*\**P<* 0.0001. n = 3. Scale bar = 200 μm.

**Supplementary Figure S4.** (A) Comparison of gene inductions among Con, KCls and MDZs groups. rPRGs: rapid PRGs, dPRGs: delayed primary response genes; SRGs: secondary response genes. **(B)** Overall changes in IGF1-related genes and pathways across these seven different groups.

**Supplementary Figure S5.** Astrocytes were also found around the tumor to restrict glioma growth in MDZ group. Tumor regions are located to the right of the dotted line. Scale bar=100μm.

**Supplementary Figure S6.** (A) Complete western blot of p-PI3K/PI3K expression in GL261 cells. **(B)** Complete western blot of p-AKT/AKT expression in GL261 cells. **(C)** Complete western blot of p-PI3K/PI3K and p-AKT/AKT expression in glioma cells in IGFBP3 experiments. **(D)** Complete western blot of p-IGF1R/IGF1R expression in GL261 cells after Linsitinib (OSI-906) treated.

## References

1 Xu S, Tang L, Li X, Fan F, Liu Z. Immunotherapy for glioma: current management and future application. Cancer letters 2020, 476: 1–12

2 Lee S, Weiss T, Bühler M, Mena J, Lottenbach Z, Wegmann R, Sun M, et al. High-throughput identification of repurposable neuroactive drugs with potent anti-glioblastoma activity. Nat Med 2024: 1–13

3 Nejo T, Krishna S, Yamamichi A, Lakshmanachetty S, Jimenez C, Lee KY, Baker DL, et al. Glioma-neuronal circuit remodeling induces regional immunosuppression. Nature Communications 2025, 16: 4770

4 Ohgaki H, Kleihues P. Population-based studies on incidence, survival rates, and genetic alterations in astrocytic and oligodendroglial gliomas. J Neuropathol Exp Neurol 2005, 64: 479–489

5 Krex D, Klink B, Hartmann C, von Deimling A, Pietsch T, Simon M, Sabel M, et al. Long-term survival with glioblastoma multiforme. Brain 2007, 130: 2596–2606

6 Yu-Ju Wu C, Chen C-H, Lin C-Y, Feng L-Y, Lin Y-C, Wei K-C, Huang C-Y, et al. CCL5 of glioma-associated microglia/macrophages regulates glioma migration and invasion via calcium-dependent matrix metalloproteinase 2. Neuro-oncology 2020, 22: 253–266

7 Chen J, Li Y, Yu T-S, McKay RM, Burns DK, Kernie SG, Parada LF. A restricted cell population propagates glioblastoma growth after chemotherapy. Nature 2012, 488: 522–526

8 Towner RA, Gillespie DL, Schwager A, Saunders DG, Smith N, Njoku CE, Krysiak RS, et al. Regression of glioma tumor growth in F98 and U87 rat glioma models by the Nitrone OKN-007. Neuro-oncology 2013, 15: 330–340

9 Hu Y, Zhou L, Wang Z, Ye Z, Liu H, Lu Y, Qi Z, et al. Assembled Embedded 3D Hydrogel System for Asynchronous Drug Delivery to Inhibit Postoperative Recurrence of Malignant Glioma and Promote Neurological Recovery. Advanced Functional Materials 2024: 2401383

10 Winkler F, Venkatesh HS, Amit M, Batchelor T, Demir IE, Deneen B, Gutmann DH, et al. Cancer neuroscience: State of the field, emerging directions. Cell 2023, 186: 1689–1707

11 Mancusi R, Monje M. The neuroscience of cancer. Nature 2023, 618: 467–479

12 Suvà ML, Rheinbay E, Gillespie SM, Patel AP, Wakimoto H, Rabkin SD, Riggi N, et al. Reconstructing and reprogramming the tumor-propagating potential of glioblastoma stem-like cells. Cell 2014, 157: 580–594

13 Lee JH, Lee JE, Kahng JY, Kim SH, Park JS, Yoon SJ, Um J-Y, et al. Human glioblastoma arises from subventricular zone cells with low-level driver mutations. Nature 2018, 560: 243–247

14 Venkatesh HS, Morishita W, Geraghty AC, Silverbush D, Gillespie SM, Arzt M, Tam LT, et al. Electrical and synaptic integration of glioma into neural circuits. Nature 2019, 573: 539–545

15 Taylor KR, Barron T, Hui A, Spitzer A, Yalçin B, Ivec AE, Geraghty AC, et al. Glioma synapses recruit mechanisms of adaptive plasticity. Nature 2023, 623: 366–374

16 Huang-Hobbs E, Cheng Y-T, Ko Y, Luna-Figueroa E, Lozzi B, Taylor KR, McDonald M, et al. Remote neuronal activity drives glioma progression through SEMA4F. Nature 2023, 619: 844–850

17 Caragher SP, Shireman JM, Huang M, Miska J, Atashi F, Baisiwala S, Park CH, et al. Activation of dopamine receptor 2 prompts transcriptomic and metabolic plasticity in glioblastoma. Journal of Neuroscience 2019, 39: 1982–1993

18 Yu K, Lin C-CJ, Hatcher A, Lozzi B, Kong K, Huang-Hobbs E, Cheng Y-T, et al. PIK3CA variants selectively initiate brain hyperactivity during gliomagenesis. Nature 2020, 578: 166–171

19 Venkatesh HS, Johung TB, Caretti V, Noll A, Tang Y, Nagaraja S, Gibson EM, et al. Neuronal activity promotes glioma growth through neuroligin-3 secretion. Cell 2015, 161: 803–816

20 Venkatesh HS, Tam LT, Woo PJ, Lennon J, Nagaraja S, Gillespie SM, Ni J, et al. Targeting neuronal activity-regulated neuroligin-3 dependency in high-grade glioma. Nature 2017, 549: 533-+

21 Chen P, Wang W, Liu R, Lyu J, Zhang L, Li B, Qiu B, et al. Olfactory sensory experience regulates gliomagenesis via neuronal IGF1. Nature 2022, 606: 550–556

22 Pan Y, Hysinger JD, Barron T, Schindler NF, Cobb O, Guo X, Yalçın B, et al. NF1 mutation drives neuronal activity-dependent initiation of optic glioma. Nature 2021, 594: 277–282

23 Johung T, Monje M. Neuronal activity in the glioma microenvironment. Curr Opin Neurobiol 2017, 47: 156–161

24 Jung E, Alfonso J, Monyer H, Wick W, Winkler F. Neuronal signatures in cancer. Int J Cancer 2020, 147: 3281–3291

25 Yang Y, Yang C, Chen X, Jiang Y, Lei X, Ma K, Quan Y, et al. Long-range cholinergic input promotes glioblastoma progression. Cancer Cell 2025, 43

26 Rzeski W, Turski L, Ikonomidou C. Glutamate antagonists limit tumor growth. Proc Natl Acad Sci U S A 2001, 98: 6372–6377

27 Pallud J, Le Van Quyen M, Bielle F, Pellegrino C, Varlet P, Cresto N, Baulac M, et al. Cortical GABAergic excitation contributes to epileptic activities around human glioma. Sci Transl Med 2014, 6: 244ra289

28 Shard C, Jones AC, Fouladzadeh A, Palethorpe HM, Francis A, Boyle Y, Ormsby RJ, et al. Novel GABAAR antagonists target networked gene hubs at the leading-edge in high-grade gliomas. Neuro Oncol 2025,

29 Barron T, Yalçın B, Su M, Byun YG, Gavish A, Shamardani K, Xu H, et al. GABAergic neuron-to-glioma synapses in diffuse midline gliomas. Nature 2025, 639: 1060–1068

30 Curry RN, Ma Q, McDonald MF, Ko Y, Srivastava S, Chin P-S, He P, et al. Integrated electrophysiological and genomic profiles of single cells reveal spiking tumor cells in human glioma. Cancer Cell 2024, 42: 1713–1728.e1716

31 Yang Y, Ren L, Li W, Zhang Y, Zhang S, Ge B, Yang H, et al. GABAergic signaling as a potential therapeutic target in cancers. Biomed Pharmacother 2023, 161: 114410

32 Buckingham SC, Campbell SL, Haas BR, Montana V, Robel S, Ogunrinu T, Sontheimer H. Glutamate release by primary brain tumors induces epileptic activity. Nat Med 2011, 17: 1269–1274

33 Anastasaki C, Mo J, Chen J-K, Chatterjee J, Pan Y, Scheaffer SM, Cobb O, et al. Neuronal hyperexcitability drives central and peripheral nervous system tumor progression in models of neurofibromatosis-1. Nature communications 2022, 13: 2785

34 Wang C, Datoo T, Zhao H, Wu L, Date A, Jiang C, Sanders RD, et al. Midazolam and Dexmedetomidine Affect Neuroglioma and Lung Carcinoma Cell Biology In Vitro and In Vivo. Anesthesiology 2018, 129: 1000–1014

35 Oshima Y, Sano M, Kajiwara I, Ichimaru Y, Itaya T, Kuramochi T, Hayashi E, et al. Midazolam exhibits antitumour and anti-inflammatory effects in a mouse model of pancreatic ductal adenocarcinoma. British Journal of Anaesthesia 2022, 128: 679–690

36 Qi Y, Yao X, Du X. Midazolam inhibits proliferation and accelerates apoptosis of hepatocellular carcinoma cells by elevating microRNA-124-3p and suppressing PIM-1. IUBMB Life 2020, 72: 452–464

37 Peng Y-R, He S, Marie H, Zeng S-Y, Ma J, Tan Z-J, Lee SY, et al. Coordinated changes in dendritic arborization and synaptic strength during neural circuit development. Neuron 2009, 61: 71–84

38 Balk M, Hentschke H, Rudolph U, Antkowiak B, Drexler B. Differential depression of neuronal network activity by midazolam and its main metabolite 1-hydroxymidazolam in cultured neocortical slices. Sci Rep 2017, 7: 3503

39 Yao M, Ventura PB, Jiang Y, Rodriguez FJ, Wang L, Perry JSA, Yang Y, et al. Astrocytic trans-Differentiation Completes a Multicellular Paracrine Feedback Loop Required for Medulloblastoma Tumor Growth. Cell 2020, 180

40 Quail DF, Bowman RL, Akkari L, Quick ML, Schuhmacher AJ, Huse JT, Holland EC, et al. The tumor microenvironment underlies acquired resistance to CSF-1R inhibition in gliomas. Science 2016, 352: aad3018

41 Lee S-C, Nakata S, Alaali L, Wang K, Tsai P-C, Pham K, Orr BA, et al. BCOR loss promotes both retinoblastoma growth and susceptibility to IGF1R inhibition. Neuro-oncology 2025, 27: 1715–1728

42 Aoki Y, Hashizume R, Ozawa T, Banerjee A, Prados M, James CD, Gupta N. An experimental xenograft mouse model of diffuse pontine glioma designed for therapeutic testing. J Neurooncol 2012, 108: 29–35

43 Ledur PF, Liu C, He H, Harris AR, Minussi DC, Zhou H-Y, Shaffrey ME, et al. Culture conditions tailored to the cell of origin are critical for maintaining native properties and tumorigenicity of glioma cells. Neuro-oncology 2016, 18: 1413–1424

44 Oshima Y, Sano M, Kajiwara I, Ichimaru Y, Itaya T, Kuramochi T, Hayashi E, et al. Midazolam exhibits antitumour and anti-inflammatory effects in a mouse model of pancreatic ductal adenocarcinoma. Br J Anaesth 2022, 128: 679–690

45 Love MI, Huber W, Anders S. Moderated estimation of fold change and dispersion for RNA-seq data with DESeq2. Genome Biology 2014, 15: 1–21

46 Kumar L, Futschik ME. Mfuzz: a software package for soft clustering of microarray data. Bioinformation 2007, 2: 5

47 Badia-i-Mompel P, Vélez Santiago J, Braunger J, Geiss C, Dimitrov D, Müller-Dott S, Taus P, et al. decoupleR: ensemble of computational methods to infer biological activities from omics data. Bioinformatics Advances 2022, 2: vbac016

48 Perry B. Shieh AG. Molecular mechanisms underlying activity-dependent regulation of BDNF expression. Journal of Neurobiology 1999,

49 Key TJ, Appleby PN, Reeves GK, Roddam AW. Insulin-like growth factor 1 (IGF1), IGF binding protein 3 (IGFBP3), and breast cancer risk: pooled individual data analysis of 17 prospective studies. Lancet Oncol 2010, 11: 530–542

50 Pollak M. Insulin and insulin-like growth factor signalling in neoplasia. Nat Rev Cancer 2008, 8: 915–928

51 Noorolyai S, Shajari N, Baghbani E, Sadreddini S, Baradaran B. The relation between PI3K/AKT signalling pathway and cancer. Gene 2019, 698: 120–128

52 Jiang N, Dai Q, Su X, Fu J, Feng X, Peng J. Role of PI3K/AKT pathway in cancer: the framework of malignant behavior. Molecular biology reports 2020, 47: 4587–4629

53 He Y, Sun MM, Zhang GG, Yang J, Chen KS, Xu WW, Li B. Targeting PI3K/Akt signal transduction for cancer therapy. Signal Transduct Target Ther 2021, 6: 425

54 Tyssowski KM, DeStefino NR, Cho J-H, Dunn CJ, Poston RG, Carty CE, Jones RD, et al. Different Neuronal Activity Patterns Induce Different Gene Expression Programs. Neuron 2018, 98

55 Faust Akl C, Andersen BM, Li Z, Giovannoni F, Diebold M, Sanmarco LM, Kilian M, et al. Glioblastoma-instructed astrocytes suppress tumour-specific T cell immunity. Nature 2025, 643: 219–229

56 Wu J, Li R, Wang J, Zhu H, Ma Y, You C, Shu K. Reactive Astrocytes in Glioma: Emerging Opportunities and Challenges. Int J Mol Sci 2025, 26

57 Venkatesh H, Monje M. Neuronal Activity in Ontogeny and Oncology. Trends Cancer 2017, 3: 89–112

58 Tian A, Kang B, Li B, Qiu B, Jiang W, Shao F, Gao Q, et al. Oncogenic State and Cell Identity Combinatorially Dictate the Susceptibility of Cells within Glioma Development Hierarchy to IGF1R Targeting. Adv Sci (Weinh) 2020, 7: 2001724

59 Li K, Duan M, Lu Q, Liu J, He M, Zhang Y. Advances in neuroscientific mechanisms and therapies for glioblastoma. iScience 2025, 28: 113347

60 Armstrong TS, Grant R, Gilbert MR, Lee JW, Norden AD. Epilepsy in glioma patients: mechanisms, management, and impact of anticonvulsant therapy. Neuro-oncology 2016, 18: 779–789

61 Drumm M, Wang W, Sears TK, Bell-Burdett K, Javier R, Cotton KY, Webb BT, et al. Postoperative risk of IDH mutant glioma-associated seizures and their potential management with IDH mutant inhibitors. The Journal of Clinical Investigation 2023,

62 Maschio M, Aguglia U, Avanzini G, Banfi P, Buttinelli C, Capovilla G, Casazza MML, et al. Management of epilepsy in brain tumors. Neurol Sci 2019, 40: 2217–2234

63 Zhang Y, Duan W, Chen L, Chen J, Xu W, Fan Q, Li S, et al. Potassium ion channel modulation at cancer-neural interface enhances neuronal excitability in epileptogenic glioblastoma multiforme. Neuron 2024,

64 Barr J, Fraser GL, Puntillo K, Ely EW, Gélinas C, Dasta JF, Davidson JE, et al. Clinical practice guidelines for the management of pain, agitation, and delirium in adult patients in the intensive care unit. Critical care medicine 2013, 41: 263–306

65 Kienitz R, Kay L, Beuchat I, Gelhard S, Von Brauchitsch S, Mann C, Lucaciu A, et al. Benzodiazepines in the Management of Seizures and Status Epilepticus: A Review of Routes of Delivery, Pharmacokinetics, Efficacy, and Tolerability. Cns Drugs 2022, 36: 951–975

66 Sun T, Chen J, Sun X, Wang G. Midazolam increases cisplatin-sensitivity in non-small cell lung cancer (NSCLC) via the miR-194-5p/HOOK3 axis. Cancer Cell Int 2021, 21: 401

