## Supplementary figures and images for "Midazolam suppresses glioma progression by attenuating neuronal activity and downregulating IGF1 signaling"

### Supplementary Fig.1

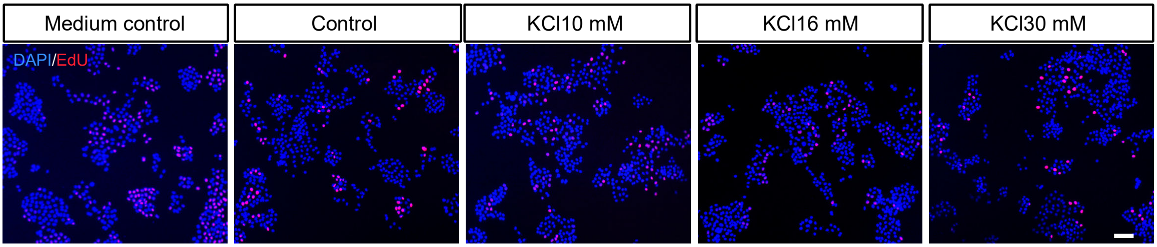

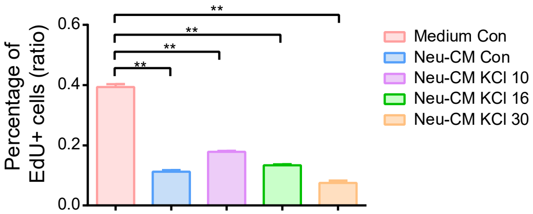

### Supplementary Fig.2

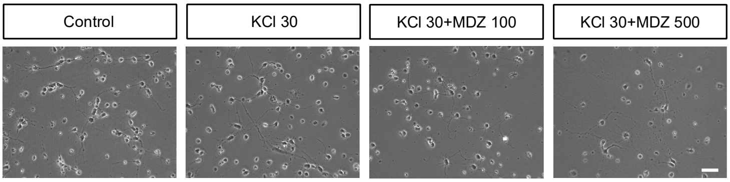

### Supplementary Fig.3

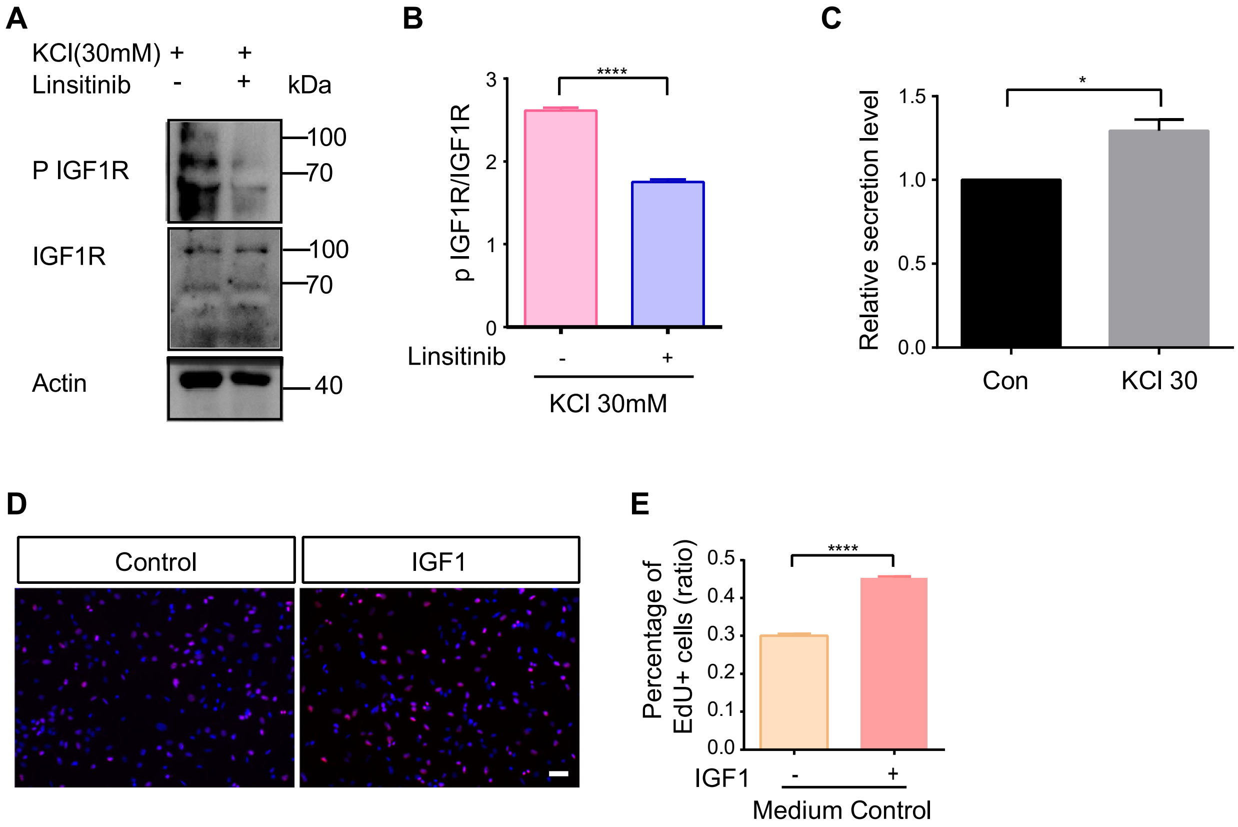

### Supplementary Fig.4

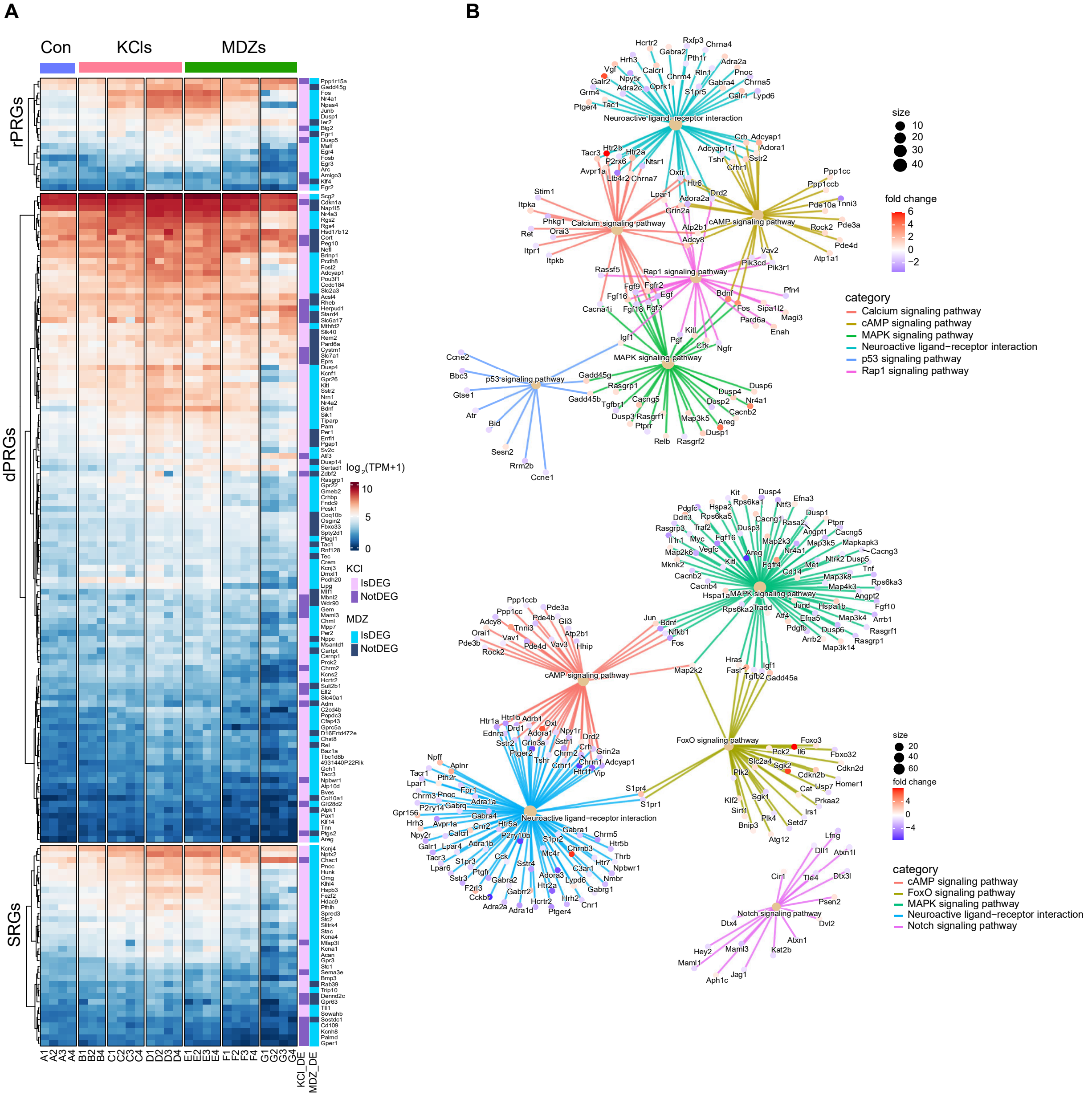

### Supplementary Fig.5

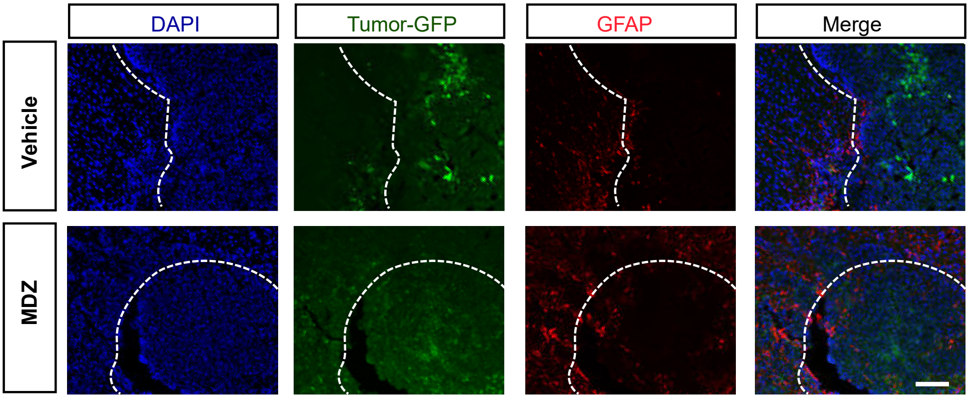
