## Supplementary Fig.6 for "Midazolam suppresses glioma progression by attenuating neuronal activity and downregulating IGF1 signaling"

A

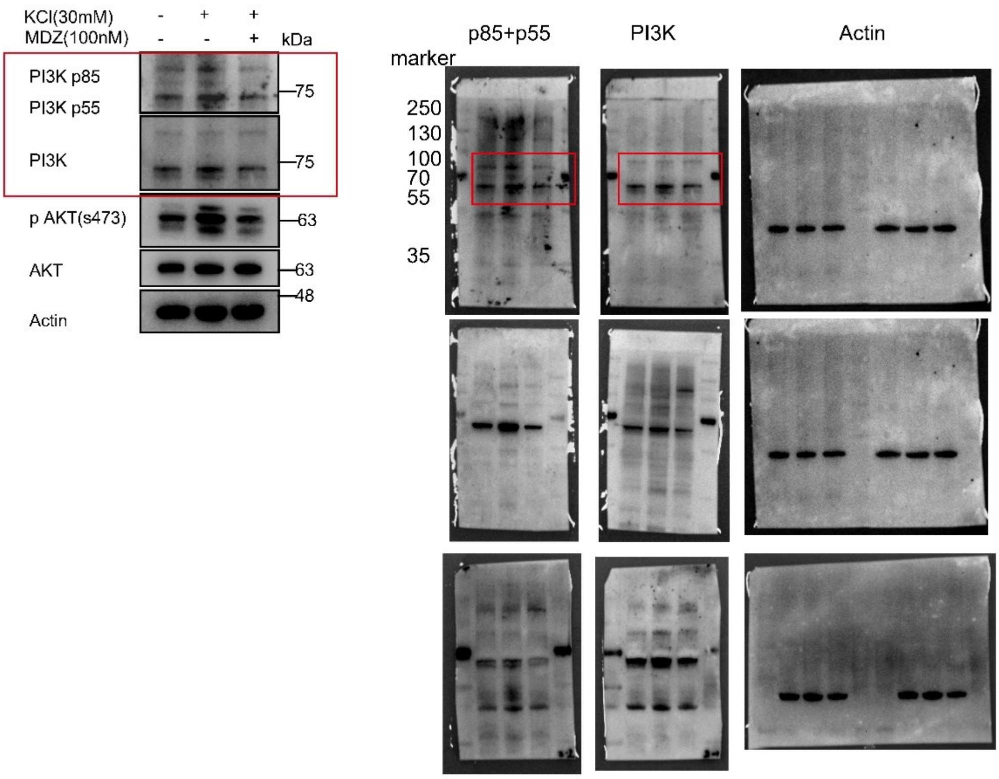
KCl(30mM) - + +
MDZ(100nM) - - +
kDa
PI3K p85
PI3K p55
75
PI3K
75
p AKT(s473)
63
AKT
63
48
Actin
p85+p55
PI3K
Actin
marker
250
130
100
70
55
35

B

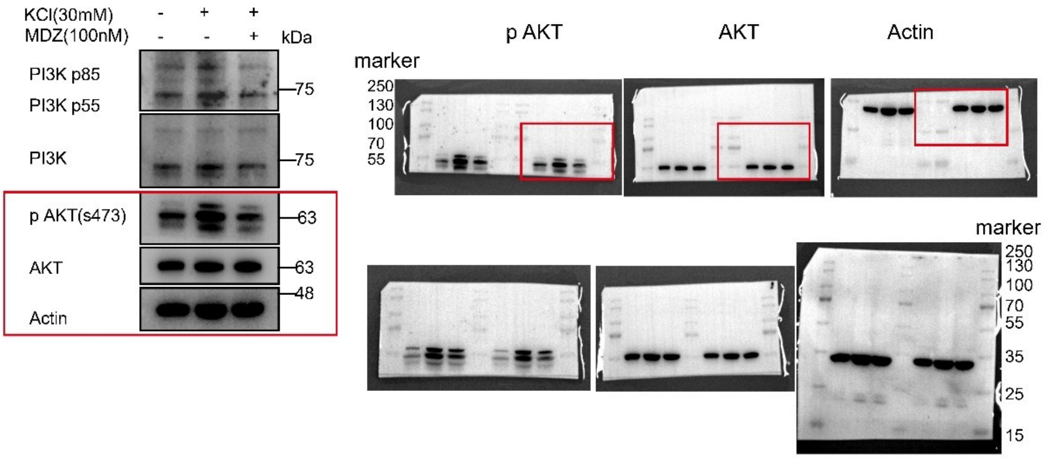
KCl(30mM) - + +
MDZ(100nM) - - +
kDa
PI3K p85
PI3K p55
75
PI3K
75
p AKT(s473)
63
AKT
63
48
Actin
p AKT
AKT
Actin
marker
250
130
100
70
55
marker
250
130
100
70
55
35
25
15

**C**

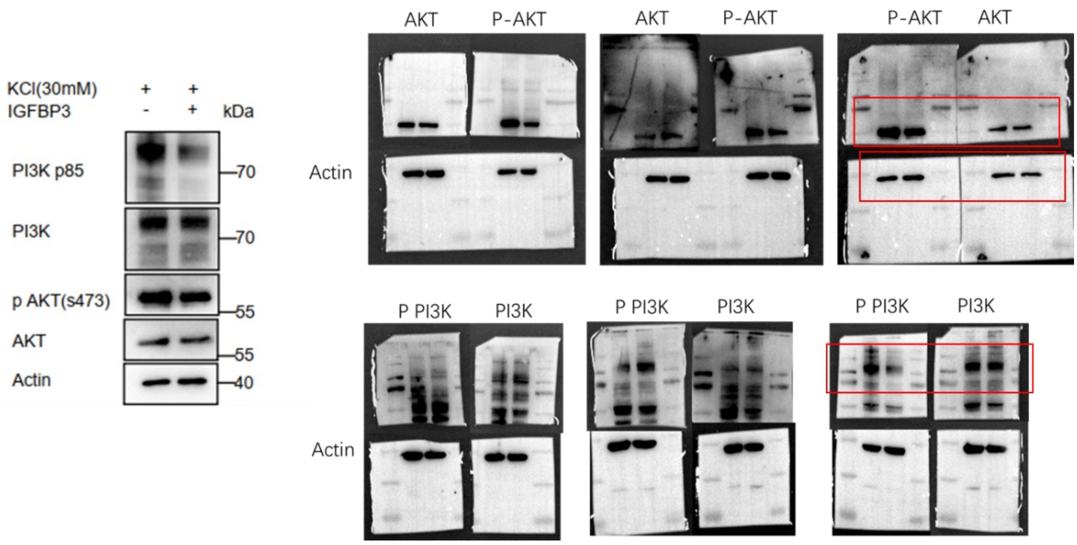


**D**

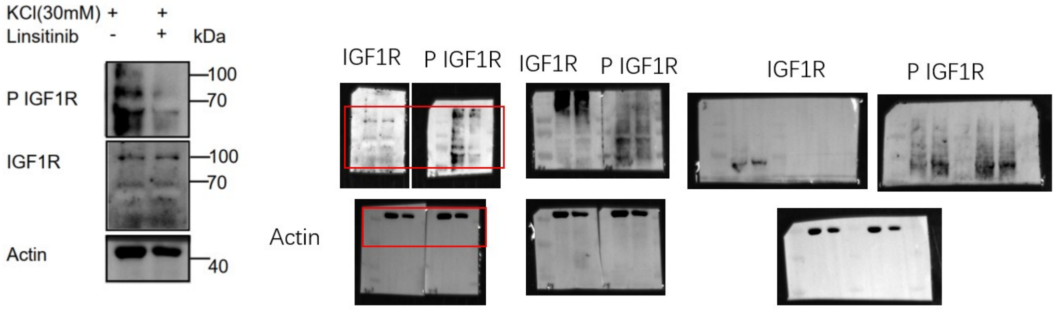


Western blotting analysis in glioma cells (GL261) cocultured with Neu-CMs. **(A)** Complete blot of p PI3K/PI3K expression in GL261 cells. **(B)** Complete blot of p AKT/AKT expression in GL261 cells. **(C)** Complete blot of p PI3K/PI3K and p AKT/AKT expression in glioma cells in IGFBP3 experiments. **(D)** Complete blot of p IGF1R/IGF1R expression in GL261 cells after Linsitinib (OSI-906) treated.
